# Genetic purging of strongly deleterious mutations underlies black-necked crane’s unusual escape from an extinction vortex

**DOI:** 10.1101/2024.04.18.590004

**Authors:** Ning Cui, Xuntao Ma, Heqi Wu, Xiaocheng Chen, Chih-Ming Hung, Lixun Zhang, Fumin Lei, Le Yang, Dao Yan, Xiaojun Yang, Feng Dong

## Abstract

Many species are undergoing rapid demographic declines, necessitating an examination of the resulting genetic impacts. The prevailing small population paradigm posits an elevated genetic load and extinction risk. However, instances of fast recovery from severe population bottlenecks suggest alternative outcomes. To investigate this issue, we performed a population genomic analysis on the black-necked crane, analyzing 42 modern and 11 historical genomes. This study revealed substantial evidence of large-effect allele purging underlying the unexpectedly rapid population recovery following an abrupt bottleneck during the 1980s. Nevertheless, forward simulations supposing a prolonged bottleneck (e.g., five generations) predicted a reversion with negative prospects, implying that rapid population recovery served as both the cause and consequence of the species escaping from an extinction vortex. These findings shed light on a potential positive microevolutionary response to current widespread population collapses and underscore the urgency of implementing active and effective conservation strategies to reverse this trend before it becomes irreversible.

## Introduction

Global biodiversity is currently experiencing an unprecedented crisis, characterized by alarming rates of species extinction [1,2], with even common species showing rapid population declines [3]. Global vertebrate populations, encompassing 4□392 species, have decreased by an average of 68% in the past five decades [4]. This decline raises a critical question about the genomic response of populations to rapid collapses. The lack of such knowledge prevents scientific evaluations of population viabilities and impedes efficient conservation policymaking.

In addition to increased inbreeding and decreased genetic diversity, the genetic consequences of population declines are particularly marked by the accumulation of deleterious mutations—genetic load [5]. In large, stable populations, genetic load is typically maintained at a relatively low level due to a balance between mutagenesis and purifying selection [6]. However, demographic contractions can disrupt this balance, exerting an enigmatic influence on genetic load [5], an area that remains contentious in the fields of evolutionary biology and conservation [6]. Nevertheless, recent studies suggest that the velocity of population decline is a critical factor [7, 8].

Gradual and prolonged declines are thought to reduce recessive detrimental alleles through intensive purifying selection—a process of genetic purging—which may facilitate long-term persistence of species, even in small populations [9-12]. In contrast, rapid and severe declines (i.e., bottlenecks) may accelerate the accumulation of deleterious mutations due to enhanced genetic drift and relaxed purifying selection [5]. This can induce inbreeding depression, resulting in a negative feedback on population size and ultimately elevating extinction risk by trapping the species in an ‘extinction vortex’, a concept central to the small population paradigm [9,13-15]. Nevertheless, there are several exceptional cases of rapid recovery from sudden and severe bottlenecks [16-18], suggesting complex genetic consequences of population declines. These cases serve as valuable resources for understanding the potential of genetic purging in rapidly declining populations, an area that remains to be empirically examined.

The black-necked crane (*Grus nigricollis*), a notable flagship bird species (Figure 1a, primarily breeds on the high-altitude Qinghai-Tibet Plateau (QTP) and winters at lower altitudes in southeast QTP as well as the western part of the Yunnan-Guizhou Plateau (Figure 1b; [19,20]). Despite its former ubiquity, the species is widely believed to have experienced a severe population decline to only 100-300 individuals during the late 20^th^ century [21,22], before rebounding to 15□000 by 2020 (Figure 2a) [22]. This represents one of the fastest recoveries observed in the animal kingdom, although little is known about the underlying genetic mechanism driving it. In this study, we assembled the first chromosome-level genome of the black-necked crane and compared population genomic characteristics before and after the bottleneck, with the specific goal of investigating the role of genetic purging. By comprehensively examining the genetic consequences in this recent bottleneck and fast recovery, this study aims to shed light on potential conservation strategies amid the current widespread and rapid declines in animal populations.

**Figure 1.**
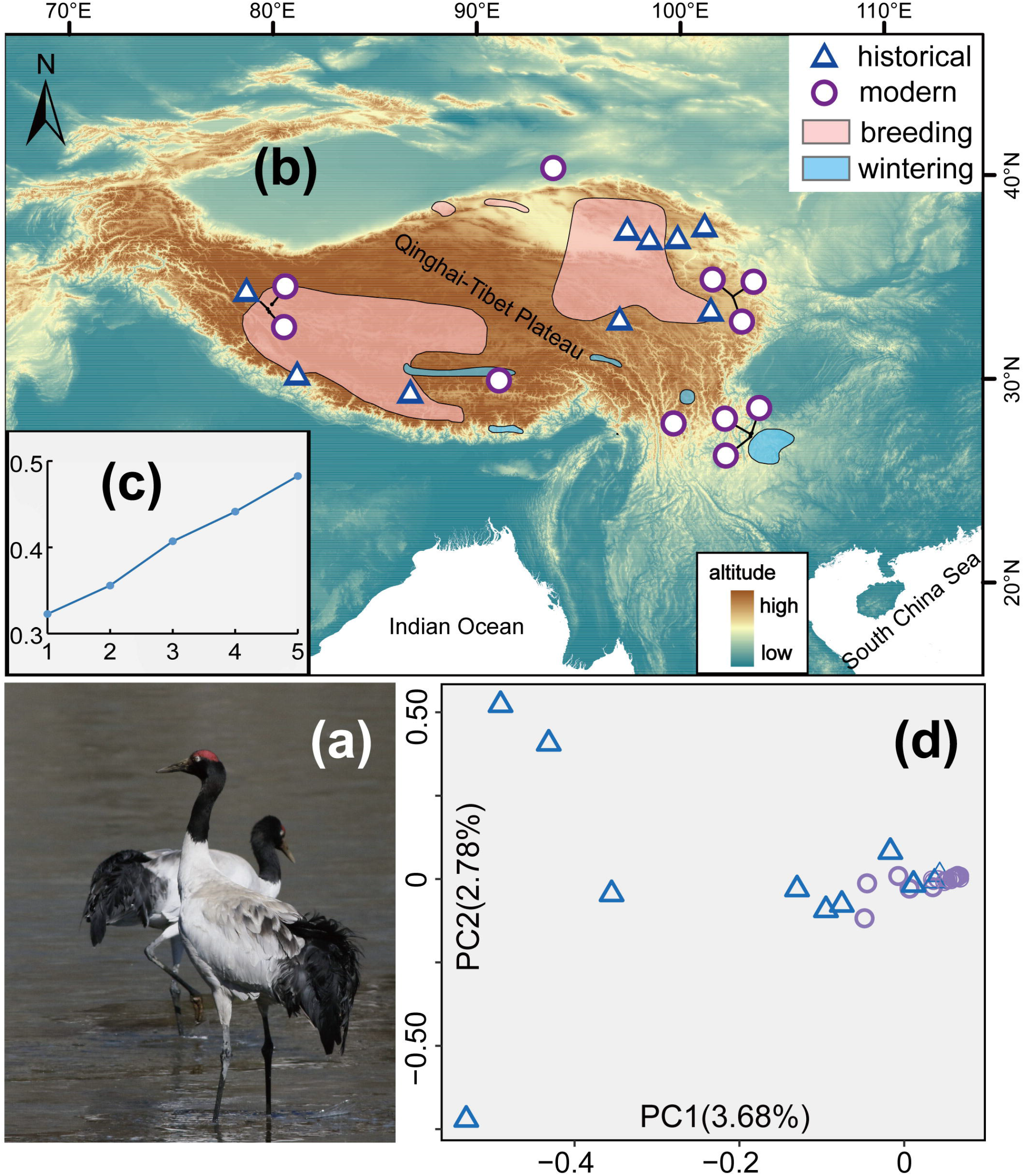
Sampling locations and population structure of black-necked crane. a: Ecological photograph (credit to Zhaxi). b: Sampling localities for historical and modern samples. c: Scatter diagrams for cross-validation (cv) errors in admixture analyses, with *K* (number of supposed genetic clusters) from 1 to 5. d: Principal component analysis (PCA) plots for all historical and modern samples.

**Figure 2.**
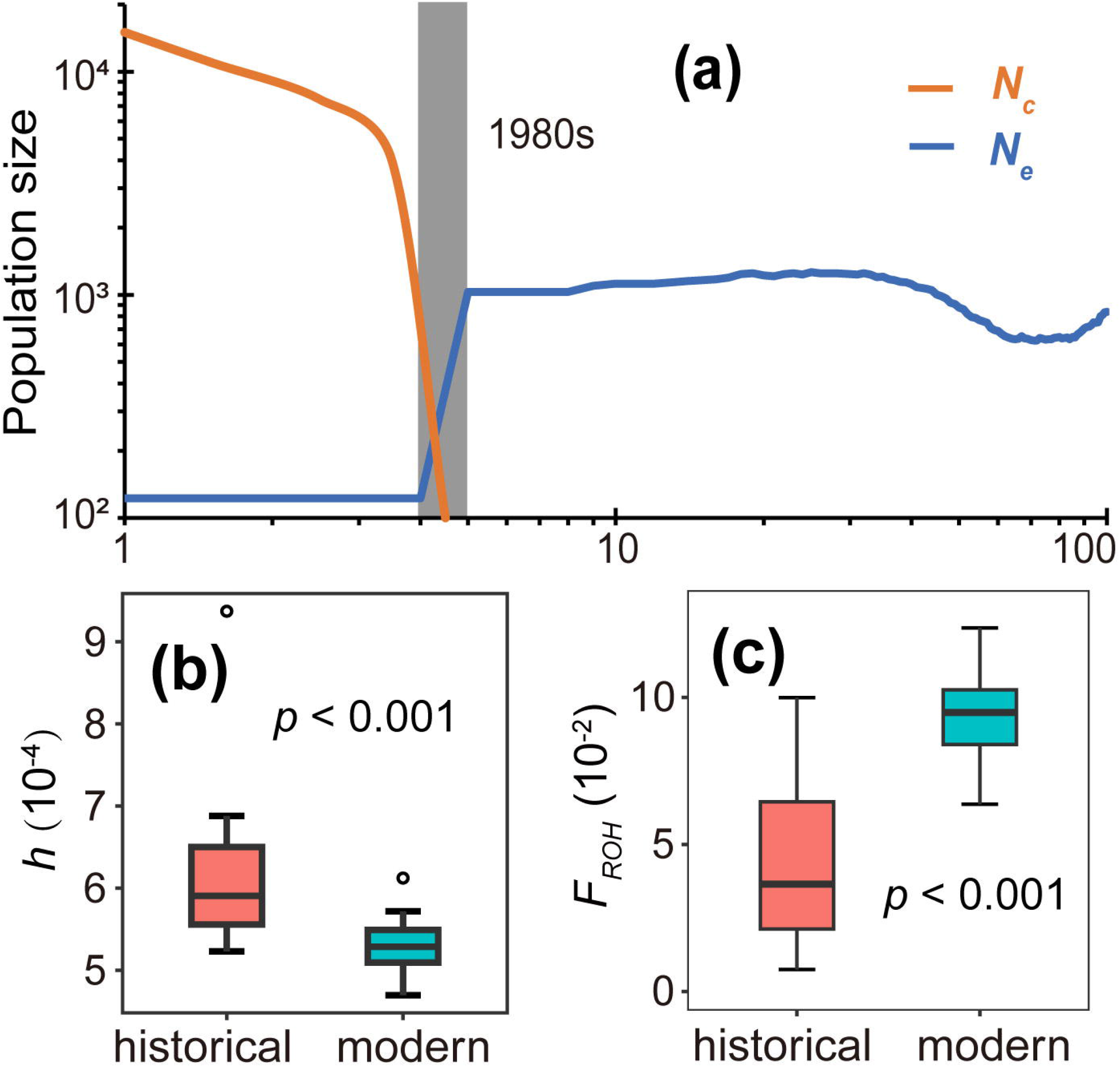
Demographic history, genomic diversity, and inbreeding estimation for black-necked crane. a: Population trajectory of recent census population size (*N*_*c*_; orange, data retrieved from ref^22^) and effective population size (*N*_*e*_; blue) with time in unit of generations. b: Boxplot of individual genome-wide heterozygosity. c: Individual inbreeding coefficients estimated as *F*_*ROH*_. For (b) and (c), horizontal lines within boxplots and bounds of boxes represent 1st and 3rd quartiles (box), median (line inside box) outliers (open circle), and 1.5× interquartile range.

## Results

### Comprehensive genomic data

We constructed a high-quality genome assembly of the black-necked crane using second- and third-generation sequencing and chromosome conformation capture-3C (Hi-C) techniques. The final assembly (Gnig-Y8) was 1.35 Gb in length and showed high continuity, with an N50 of 85.88 Mb. Notably, the top 34 largest scaffolds, out of a total of 314 scaffolds (> 4 Mbp; hereafter: pseudochromosomes) accounted for 96% of the genome (1.29 Gb), showing remarkable improvement over the previous genome version [23]. Further analysis confirmed high genomic completeness of this assembly, achieving a retrieval rate of 97% for complete Benchmarking Universal Single-Copy Orthologs (BUSCO) among birds (Supplementary Figure S1). The annotation process identified 20□963 gene models with 922□165 exons.

We resequenced a total of 52 genomes with an average coverage of 30.2x (15.9-83.8x; Supplementary Table S1) and compiled four comprehensive population genomic datasets for subsequent analyses. The comparative genomic dataset comprised 2□829□286 high-quality autosomal biallelic single nucleotide polymorphisms (SNPs) derived from 41 modern black-necked cranes spanning their extant distribution range and 11 historical samples predating the recent bottleneck (Figure 1b), which were used to estimate genomic diversity and genetic load, while a subset of 560□083 putatively unlinked SNPs was used for genetic clustering analysis. We also compiled a dataset of 1□319□028 automatic biallelic SNPs from modern samples to reconstruct recent demography and a subset of 21□590 putatively unlinked SNPs to estimate contemporary effective population size (*N*_*e*_).

### Lack of population genetic structure

We conducted admixture and principal component analysis (PCA) of historical and modern black-necked cranes to examine potential genetic stratification. Admixture analysis indicated that the lowest cross-validation (CV) error occurred at *K (number of supposed genetic clusters)* =1 (Figure 1), suggesting a single breeding population origin for all samples. The PCA results further demonstrated markedly reduced heterogeneity in modern genomes compared to historical genomes (Figure 1d).

### Recent sharp population bottleneck

We retrieved details on recent population trajectories for the black-necked crane using the optimal GONE algorithm [24, 25], which uses a linkage disequilibrium spectrum and provides high resolution within the last 100 generations (1□000 years here) [24]. Results revealed a sharp bottleneck during the 1980s, with an eight-fold *N*_*e*_ decline from approximately 1□000 to 122 diploids (Figure 2a). Similarly, independent estimation using NeEstimator [4] based on putatively unlinked loci suggested a contemporary *N*_*e*_ of 127 diploids (95% confidence interval: 112 to 137).

### Decreased genomic diversity and increased inbreeding

To characterize genomic diversity before and after the bottleneck, we first calculated genome-wide genetic diversity (π), which was 1.2 times higher in the historical than in the modern genomes (Wilcoxon signed rank test, *P*<0.001; Supplementary Figure S2). Second, we estimated genome-wide heterogeneity (*h*; average likelihood of heterozygosity for each site), which was 1.1 times higher in the historical than in the modern genomes (Wilcoxon rank sum test, *P*<0.001; Figure 2b). Third, we measured inbreeding coefficients (*F*_*ROH*_) as the proportion of the genome covered by runs of homogeneity (ROHs), finding a 2.6-fold increase in modern genomes compared to historical genomes (Wilcoxon signed rank test, *P*<0.001; Figure 2c).

### Increased genetic load in homozygous state

We measured and compared genetic load in historical and modern genomes using three distinct approaches. First, we estimated genetic load by counting the number of derived deleterious variants in coding regions and classified them into three types: those in heterozygotes as masked genetic load, those in homozygotes as realized genetic load, and the combined total of the first two types as total genetic load. We categorized the potential impacts of genetic load as High and Moderate using SnpEff [26] and compared their distributions between historical and modern genomes (see Methods and Materials). In the High-impact category, we observed a 22.6% reduction in total genetic load in the modern genomes relative to historical ones (Wilcoxon rank sum test, *P*<0.001; Figure 3a), with masked genetic load showing a 3.2% reduction (Welch’s two-sample *t*-test, *P*<0.001; Figure 3b) but realized genetic load showing a 26.6% increase (Welch’s two-sample *t*-test, *P*<0.001; Figure 3c). In the Moderate impact category, we found no difference in total genetic load (Wilcoxon rank sum test, *P*=0.053; Supplementary Figure S3a), but a 2.4% decrease in masked genetic load (Welch’s two-sample *t*-test, *P*<0.001; Supplementary Figure S3b) and a 21.5% increase in realized genetic load in the modern genomes compared with the historical genomes (Welch’s two-sample *t*-test, *P*<0.001; Supplementary Figure S3c).

**Figure 3.**
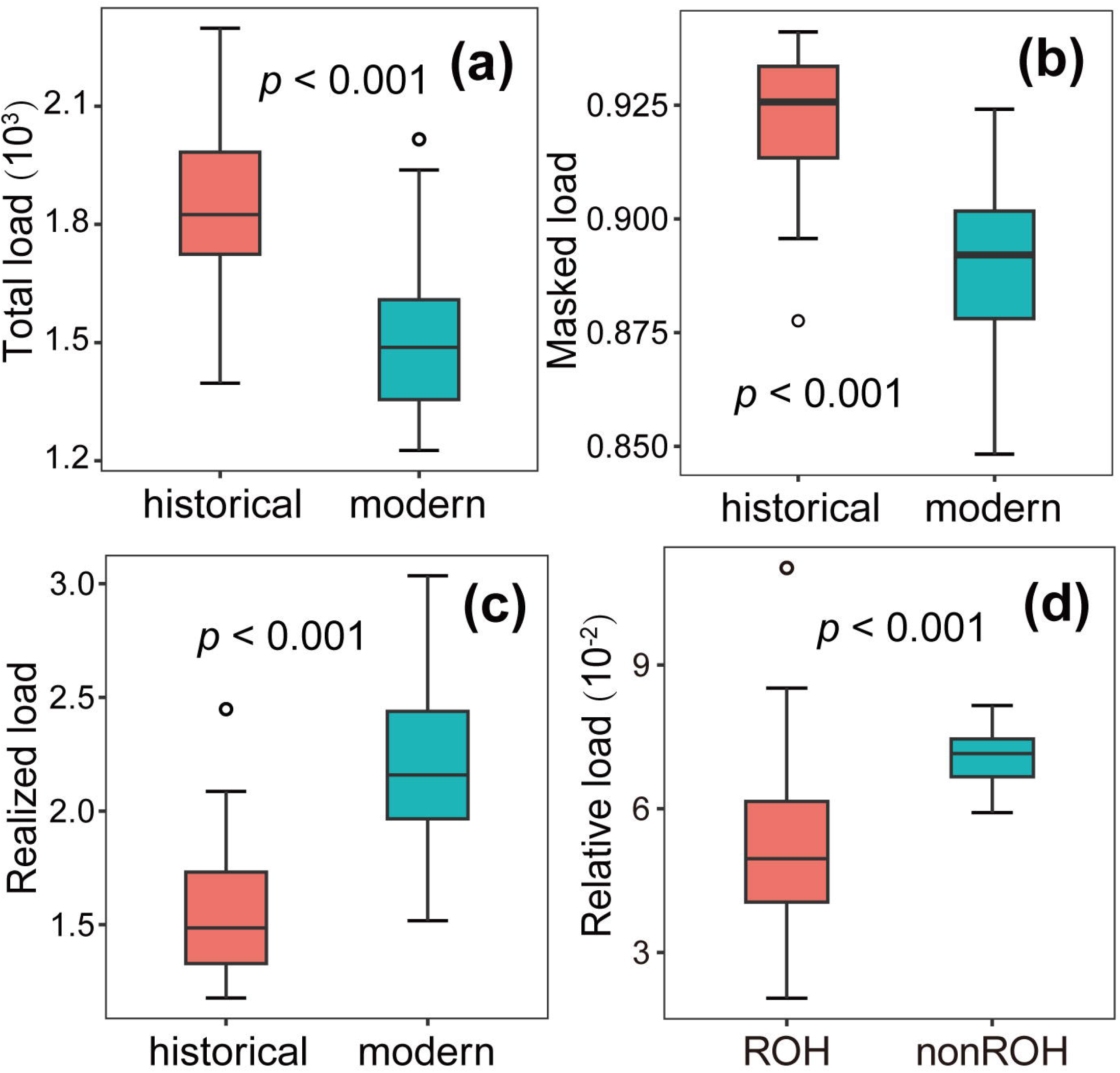
Genetic load estimation on coding regions for black-necked crane. A: Total genetic load; B: Masked genetic load; C: Realized genetic load; and D: Relative genetic load inside and outside ROHs based on High-impact category from SnpEff analysis. Boxplot shows 1st and 3rd quartiles (box), median (line inside box), outliers (open circle), and 1.5× interquartile range.

Second, we estimated and compared the relative frequency of derived alleles (R_modern/historical_) for the High- and Moderate-impact variants in the historical and modern genomes. The High-impact category showed an excess of deleterious derived alleles (R_modern/historical_=1.24; Supplementary Figure S4) in homozygotes and a deficit in heterozygotes (R_modern/historical_=0.93; Supplementary Figure S4) in the modern genomes, indicating higher realized genetic load and lower masked genetic load. The Moderate-impact category showed a congruent excess in both homozygotes and heterozygotes (R_modern/historical_=1.23 and 1.04, respectively; Figure 4).

**Figure 4.**
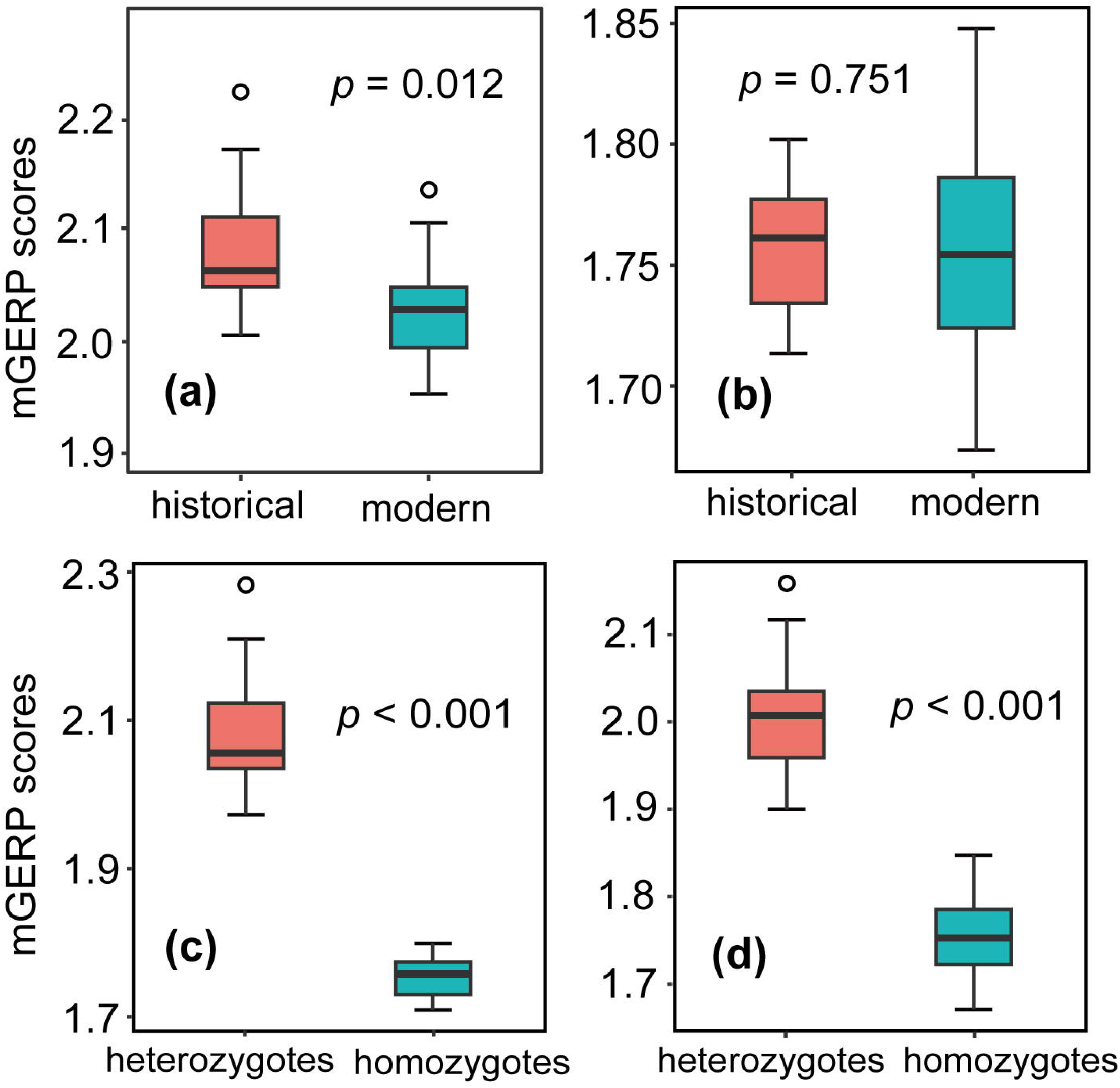
Selection coefficient estimation based on mGERP scores for the black-necked crane. Comparisons were conducted for (A) heterozygotes and (B) homozygotes between historical and modern genomes, as well as for individual genomes among (C) historical and (D) modern genomes. Boxplot comprises 1st and 3rd quartiles (box), median (line inside box), outliers (open circle), and 1.5× interquartile range.

Third, we compared the relative abundance of homozygous High-impact deleterious variants within and outside ROHs in the modern genomes. We found a 40.2% lower presence of these variants within ROHs (Welch’s pairwise *t*-test, *P*<0.001; Figure 3d). Given that deleterious variants within ROHs may reduce individual fitness more markedly than those outside ROHs [5], these results suggest a signature of genetic purging in the modern population.

### Constant purifying selection over the bottleneck

We examined purifying selection signals between the historical and modern genomes. First, we estimated genomic evolutionary rate profiling (GERP) scores for each site of the genomes, where loci with higher values were assumed to be under higher purifying selection pressure. We examined sites under the strongest evolutionary constraint (i.e., those with the top 100 GERP scores) and referred to their scores as the maximum GERP (mGERP) scores. We found lower mGERP scores in the modern genomes compared to the historical genomes in heterozygotes (Welch’s two-sample *t*-test, *P*=0.011; Figure 4a), but no significant difference in homozygotes (Welch’s two-sample *t*-test, *P*=0.751; Figure 4b). In addition, we found significantly higher mGERP scores in heterozygotes than homozygotes in both historical and modern genomes (Welch’s pairwise *t*-test, *P*<0.001 for both; Figure 4c-d).

Second, we conducted forward simulations and compared the individual-based selection coefficients of (partially) recessive deleterious mutations in homozygous sites between the historical and modern genomes (e.g., dominance coefficient (*h*)=0 for fully recessive and *h*=0.25 for partially recessive), a major genetic basis for inbreeding depression [8]. Given that deleterious mutations present negative selection coefficients, lower values indicate more relaxed purifying selection. We assumed a pre-bottleneck consensus population size (*N*_*c*_) of approximately 10□000 individuals given an empirical *N*_*e*_*/N*_*c*_ ratio of 0.1 [27] and a pre-bottleneck *N*_*e*_ of approximately 1□000 (Figure 2a). Recapitulation of the demographic history of the black-necked crane indicated no significant temporal difference in the minimum selection coefficient for both fully and partially recessive deleterious mutations (Wilcoxon rank sum test, *P*=0.331 and *P*=0.256, respectively; Figure 5a-b). Assuming the bottleneck lasted for five generations (approximately 50 years), we found that purifying selection was significantly relaxed for both types of recessive deleterious mutations (Wilcoxon rank sum test, *P*<0.001; Figure 5c-d).

**Figure 5.**
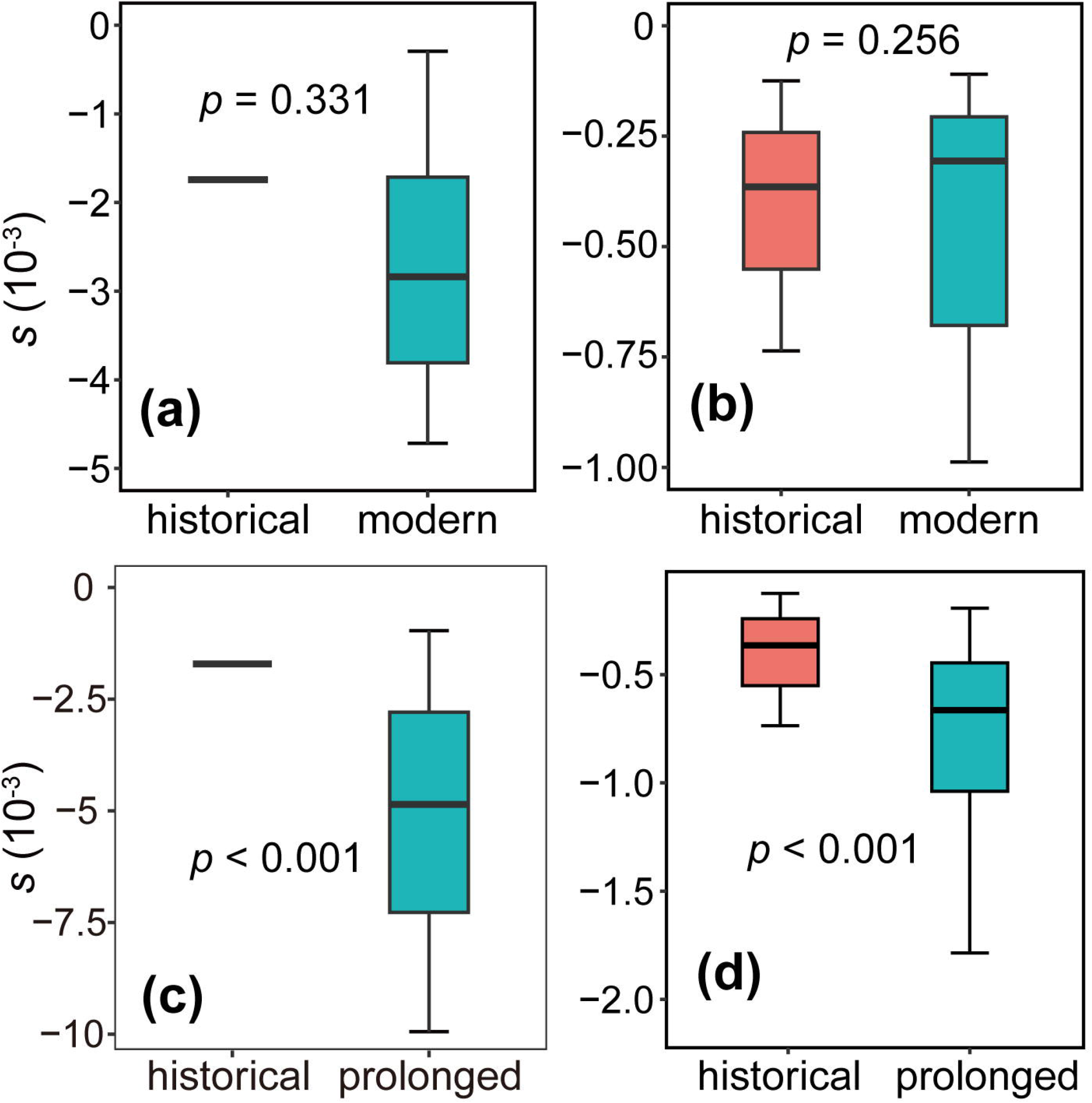
Forward simulations on selection coefficient (*s*) for deleterious mutations in homozygotes of black-necked crane. Simulations recapitulated demographic history (A-B) and supposed a prolonged bottleneck (C-D) under fully recessive (A and C) and partially recessive (B and D). Boxplot comprises 1st and 3rd quartiles (box), median (line inside box), and 1.5× interquartile range.

## Discussion

Based on comprehensive population genomic data, we identified an abrupt eight-fold reduction in the effective population size of the black-necked crane during the 1980s (Figure 2a). This decline corresponds with historical records of a severe bottleneck, which reduced the population to only 100-300 individuals, during the same period [21,22]. While this decline was initially considered a potential artifact of inadequate investigation, subsequent intensive field surveys challenged this view, showing a continual and rapid population expansion

(Figure 1a; [22]), indicating recovery from a preceding bottleneck. Support for the bottleneck event is further provided by our observations of reduced genetic heterogeneity and elevated inbreeding in modern genomes compared to historical genomes (Figure 1d and Figure 2c). Despite asymmetric sample sizes between historical and modern genomes (e.g., 11 versus 41), this sampling scheme is unlikely to introduce overt bias as both groups contained more than the minimum sample size of 4-6 individuals required for accurate estimations of genomic characteristics [28,29]. Our findings suggest that the black-necked crane remains at high genetic risk, with a contemporary *N*_*e*_ (e.g., 127) well below the generally accepted safe genetic threshold (e.g., 500; [30]), suggesting potential vulnerability to changing environments [30]. These results stand in marked contrast to the recent removal of this species from the threatened category of the IUCN Red List in 2020 [31].

According to the small population paradigm, the recent and sudden bottleneck likely trapped the black-necked crane in an extinction vortex by accelerating the accumulation of genetic load [13]. Indeed, we observed a post-bottleneck increase in realized genetic load (homozygous sites) (Figure 3c) alongside a decrease in the masked genetic load (heterozygous sites) (Figure 3b). This shift is expected as increased homozygosity, facilitated by inbreeding, converts masked genetic load into realized genetic load [13]. Moreover, this conversion is anticipated to release harmful variants hidden in heterogeneity in the historical genomes, thereby intensifying the deleterious nature of realized genetic load in the modern genomes and elevating the risk of inbreeding depression and extinction (Figure 4c-d).

In sharp contrast, the black-necked crane has demonstrated a rapid recovery following the bottleneck, with population numbers rebounding to 15□000 individuals by 2020 [22], indicating limited inbreeding depression. We propose that the rapid purging of strongly deleterious mutations likely drove this unexpected population trajectory. Notably, our observations revealed that homozygous High-impact alleles were proportionally less enriched inside than outside ROHs in the modern genomes (Figure 3d), consistent with genetic purging of strongly deleterious mutations during inbreeding. Furthermore, our empirical data (Figure 4b) and forward simulations (Figure 5a-b) demonstrated a temporally consistent level of purifying selection on homozygotes pre- and post-bottleneck. This likely acted as a filter to purge alleles with large recessive deleterious effects on fitness, a decisive source of inbreeding depression [8] due to increased exposure to homozygotes. The purging of strongly deleterious alleles by efficient purifying selection may have enabled the black-necked crane to escape an expected extinction vortex, a mechanism that may also apply to other organisms with similar rapid demographic recoveries [16,17].

Based on comprehensive examination of the potential genetic load dynamics of a sudden and severe population collapse followed by an unexpectedly fast recovery, our study sheds light on the genetic consequences of contemporary widespread and rapid population declines. Our findings stand in contrast to the existing small population paradigm by demonstrating an escape from a potential extinction vortex following drastic population decline. Nevertheless, consistent with theoretical predictions [5], forward simulations suggested that the duration of population bottlenecks may alter this genetic consequence. For example, a prolonged bottleneck lasting for five generations may lead to relaxed purifying selection and the accumulation of strongly deleterious mutations, thereby elevating the risk of inbreeding depression and extinction (Figure 5d-e). Thus, rapid population recovery may serve as both the cause and consequence of the unusual escape of the black-necked crane from an extinction vortex. Overall, our study raises hope for the prospects of current widespread and rapid population declines, while urging the need for active and effective conservation measures to reverse the trend before it is too late.

## Methods and Materials

### Sampling and sequencing

We collected blood tissue from a rescued female black-necked crane in Yunnan Province, China. We extracted high molecular weight genomic DNA using a QIAGEN® Genomic Kit (Cat#13343, QIAGEN) for shotgun short-read and PacBio long-read sequencing. For shotgun short-read data, we generated 151 bp paired-end reads from a 350 bp insert-size library using the MGISEQ-T7 platform and obtained 47.13 Gb of clean reads after quality control using fastp v0.12.6 (-n 0 -f 5 -F 5 -t 5 -T 5; [32]). For the PacBio long-read dataset, we sheared the gross genomic DNA by g-TUBEs (Covaris, USA) and prepared SMRTbell libraries with an average insert-size of 14.85 kb according to the standard PacBio protocols (Pacific Biosciences, CA, USA). Raw polymerase reads were produced using the PacBio Sequel□platform with a Sequel II Sequencing Kit 2.0 on Nextomics, resulting in 30.58 Gb of PacBio subreads after cleaning raw reads with ccs (-min-passes 1 -min-rq 0.99 -min-length 100; [33]).

To assemble hybrid genomes into chromosomes, we generated Hi-C data for BioNano mapping construction. The blood tissue was fixed through a 10 min incubation of the cross-linking reaction by vacuum infiltration in a nuclear isolation buffer supplemented with 2% formaldehyde. We terminated the crosslinking process with glycine and constructed a library using 100 units of DpnII restriction endonuclease (GATC), followed by a sequencing procedure with the same protocol as the second-generation sequencing data. Finally, we obtained 76.21 Gb of indexed Hi-C clean reads after filtering using HiC-Pro v2.8.1 [34] and fastp.

### Genome assembly

We assembled a chromosome-level reference genome using the above three complementary datasets. We initially assembled the PacBio clean reads into contigs using Hifiasm (https://github.com/chhylp123/hifiasm), against which we mapped all PacBio clean reads with BLASR [35] to correct potential sequencing errors. We then polished these contigs with shotgun clean data using NextPolish (four iterations; [36]) with default parameters. To further improve the accuracy of the initial assembly, we clustered, ordered, and oriented contigs onto chromosomes using LACHESIS [37] with Hi-C clean data (-cluster_min_re_sites=100, -cluster_max_link_density=2.5, - cluster_noninformative_rative=1.4, -order_min_n_res_in_trunk=60, - order_min_n_res_in_shreads=60). We assessed the completeness of the final assembly (Gnig-Y8) by searching 8□338 single-copy orthologs in the aves_odb10 dataset with BUSCO v4.0.5 [38]. We preliminarily obtained the annotations by mapping genes of the well-annotated chicken genome (GenBank assembly accession: GCA_000002315.5, hereafter: chicken genome) onto Gnig-Y8 using Liftoff v1.6.1 [39] and then removed gene models with in-frame STOP codons using gffread utility (-V option) from cufflinks v2.2.1 [40].

### Resequencing and variant calling

The population genomic data involved 41 modern individuals sampled from 2010 to 2020 as well as 11 historical specimens collected from 1956 to 1976, a period predating the recent bottleneck during the mid-1980s (see details in Supplementary Table S2). We resequenced these samples with a 350 bp insert-size library and 151 bp read length, removed adapters with Cutadapt v2.10 [41], masked low-quality reads (e.g., Phred score<15) with Seqtk [42], and performed the mapping procedure (including alignment, sorting, and deduplication) using Burrows-Wheeler Aligner v0.7.17 [43], SAMtools v1.7 [44], and Genome Analysis Tool Kit (GATK) v4.0.3 [45]. We identified raw variants using Sambamba v0.6.6 [46] and implemented various filtering procedures for downstream analyses.

### Genetic clustering analysis

We conducted genetic clustering analysis using the likelihood-based algorithm in Admixture v1.3.0 (with ten runs of each *K=*1 to 5) [47] and PCA in GCTA v1.92.2beta [48]. To prepare the inputs, we extracted autosomal biallelic SNPs with VCFtools v0.1.17 (‘-f Qual=20/MinMQ=30’ & ‘-min-alleles 2 -max-alleles 2 -remove-indels -mac 2 -max-missing 1.0’; [49]), then filtered for potential linkage disequilibrium using PLINK v1.0.7 (‘-indep 50 5 2’; [50]), finally obtaining 560□083 putatively unlinked SNPs.

### Demographic reconstruction

We estimated the recent population history of the black-necked crane using GONE [51] based on linkage disequilibrium data from modern samples. To scale demographic events in calendar time, we assumed a generation length of 10 years [52] and estimated there combination rate using the following procedure. We first calculated the population recombination rate (ρ=4*N*_*e*_*r*□, where *r* is the recombination rate per generation) with a 50 kb nonoverlapping window using the R package FastEPRR [53]. We prepared inputs by filtering the raw variants from Sambamba using VCFtools with the following parameters: ‘-f Qual=20/MinMQ=30’ & ‘-max-missing 1.0 -minDP 5 -min-alleles 2 -max-alleles 2 -remove-indels -maf 0.05’, finally obtaining 1□319□028 automatic biallelic SNPs. We also calculated the population mutation rate (θ=4*N*_*e*_μ, where μ is the recombination rate per generation) using ANGSD v0.928 (‘-doSaf 1 -remove_bads -only_proper_pairs 1 -minMapQ 30 -minQ 20 - minInd 41 | realSFS | thetaStat’; [54]) with the same window size based on the mapped BAM files. We finally estimated the recombination rate as 3.42×10^-8^ using the equation *r*=(ρ*/*θ)×μ, assuming an μ of 1.45×10^-8^ per site per generation [55].

We also estimated contemporary *N*_*e*_ using a linkage disequilibrium method in NeEstimator [56]. To address the reduced precision associated with large and linked loci, we thinned the dataset previously used in the GONE analysis to a minimum distance of 50 kb, yielding a set of 21□590 SNPs.

### Genomic diversity and inbreeding estimation

We compared genetic diversity between the historical and modern samples using nucleotide diversity (π) and heterogeneity (*h*). We performed both analyses based on 2□829□286 biallelic data points, excluding singletons, with the VCFtools command: -max-missing 1.0 -mac 2 -minDP 3 -min-alleles 2 -max-alleles 2 -remove-indels. We calculated genome-wide π using VCFtools with a 50 kb window and 20 kb steps. We counted heterogeneous sites for each individual and measured the *h* index by dividing the counts by the autosomal genome length (1.17 Gb).

To assess the level of inbreeding in the black-necked crane, we calculated the length and number of ROHs for modern and historical genomes with PLINK with the following parameters: “-homozyg-window-snp 50 -homozyg-snp 50 -homozyg-window-missing 3 - homozyg-kb 100 -homozyg-density 1000”. We calculated the inbreeding coefficient (*F*_*ROH*_) for each individual by dividing the sum length of ROHs by the autosomal genome length (1.17 Gb). Higher *F*_*ROH*_ values are indicative of higher inbreeding levels.

### Genetic load calculation

We estimated genetic load dynamics between the modern and historical genomes based on functional effects of SNPs in coding regions and conservation scores under strict evolutionary constraints. Both analyses were based on the same dataset as in the *Genomic diversity and inbreeding estimation* section, with ancestral states polarized by genomic data of the hooded crane (*Grus monachus*) (GenBank assembly accession: GCA_012487855.1), the sister species of the black-necked crane [57].

First, we built a database based on the Gnig-Y8 annotation and categorized the effects of coding variants into four impact levels using SnpEff v4.3.68 [26]: a) High: variants assumed to have disruptive impact on the protein (e.g., loss-of-function and frameshift variants); b) Moderate: nondisruptive variants that may change protein effectiveness (e.g., inframe_deletion); c) Low: variants mostly harmless or unlikely to change protein behavior (e.g., synonymous variants); and d) Modifier: usually noncoding variants or variants affecting noncoding genes (e.g., downstream variants). We counted the number of total variants and respective proportions of homo- and heterozygotes (multiplied by 2 in homozygous positions) in the High and Moderate categories and compared the total, realized, and masked genetic loads between the historical and modern genomes.

Second, we estimated the relative risk of deleterious burden, R_modern/historical_ [58], based on the derived allele frequency of deleterious mutations from the historical and modern genomes. We conducted this estimation for the realized, masked, and total genetic loads using a script brrAB developed by Zheng et al. (2023) [59]. Here, R _modern/historical_ equal to 1 corresponds to no change in frequency between historical and modern genomes, while R _modern/historical_<1 or >1 indicates an increase or a decrease in frequency in historical genomes relative to modern genomes, respectively. The distributions in the R _modern/historical_ variance were estimated based on jack-knifing across chromosomes.

Third, to test the hypothesis that deleterious variants in ROH tracts may reduce individual fitness with additive effects [5], where purging, if any, is expected to be more common than in the rest of genome, we compared the ratio of High-impact (obtained from SnpEff output) to all homozygous loci in ROH and non-ROH portions of each modern genome.

### Selection coefficient estimation

We employed three approaches to compare purifying selection coefficients between historical and modern genomes. First, we estimated the coefficients for each individual using conservation scores. In brief, we downloaded GERP scores computed for 27 sauropsids by multiple whole□genome alignment [60] and converted chicken-genome-focused scores to the black-necked crane genome using the LiftOver program (http://genome.ucsc.edu/cgi-bin/hgLiftOver). We then compared the top 100 GERP (mGERP) scores in homo- and heterozygotes between the historical and modern genomes to detect dynamics of purifying selection coefficients over the bottleneck.

Second, we conducted forward simulations using SLiM v2.0.1 [61]. In brief, we modeled individuals with a diploid genome of 2 Mb, composed of 2□000 genes, each 1 kb in length, across 34 pseudochromosomes mirroring the relative proportions and positions observed in the Gnig-Y8 assembly, assuming: a) a per-generation mutation rate of 1.45×10^-8^ [55]; b) a recombination rate of 3.42×10^-8^, as estimated in the previous *Demographic reconstruction* section between genes and no intragenic recombination; c) a relative occurrence ratio of 7:3 for neutral and deleterious mutations; and d) a gamma distribution of fitness (mean=0.03, standard deviation (sd)=0.1) used as selection coefficients of deleterious mutations [62]. We simulated neutral and deleterious variation following a 100□000 generation burn-in period, with three distinct scenarios: a) a pre-bottleneck population with an *N*_*c*_ of 10□000 individuals given an empirical *N*_*e*_*/N*_*c*_ ratio of 0.1 and a pre-bottleneck *N*_*e*_ level of approximately 1□000; b) a post-bottleneck population recapitulating the demographic history of the black-necked crane, decreasing from *N*_*c*_=10□000 to 100 individuals in a generation and rebounding immediately to 15□000 individuals over four generations; and c) a scenario similar to the second, but with a prolonged bottleneck (e.g., with 100 individuals) lasting five generations. For each scenario, we performed a range of dominance coefficients, including entirely recessive (*h*=0), partially recessive (*h*=0.25), and entirely additive (*h*=0.5). We randomly subsampled 50 individuals from the last simulation step and compared their mFEs of deleterious mutations for homo- and heterozygous mutations.

## Data availability

The scripts used in the paper were deposited in the Science Data Bank (doi:10.57760/sciencedb.10054). Genomic data were archived in the Genome Warehouse (accession number: GWHDTVR00000000) and Genome Sequence Archive (accession numbers: CRA012191 and CRA012234).

## Acknowledgments

The authors would like to thank Dr. Qiang Liu, Dr. Dejun Kong, Dr. Kai Wang, Dr. Peng He, Mr. Weixiong Luo, and Ms. Mengyin An for efforts in collecting DNA samples. This work was supported by the National Key R&D Program of China (2022YFC2601200) and Second Tibetan Plateau Scientific Expedition and Research (STEP) program (2019QZKK0501)

## Author contributions

F.D. conceived the ideas; N.C., X.T.M., F.D., and H.Q.W conducted the analyses; X.C.H., H.Q.W., L.X.Z., F.M.L., L.Y., D.Y., and X.J.Y. collected DNA samples; F.D. led the writing, with substantial input from C.M.H., which was approved by all other authors.

