## Supplementary Figures and Tables for "Genetic purging of strongly deleterious mutations underlies black-necked crane’s unusual escape from an extinction vortex"

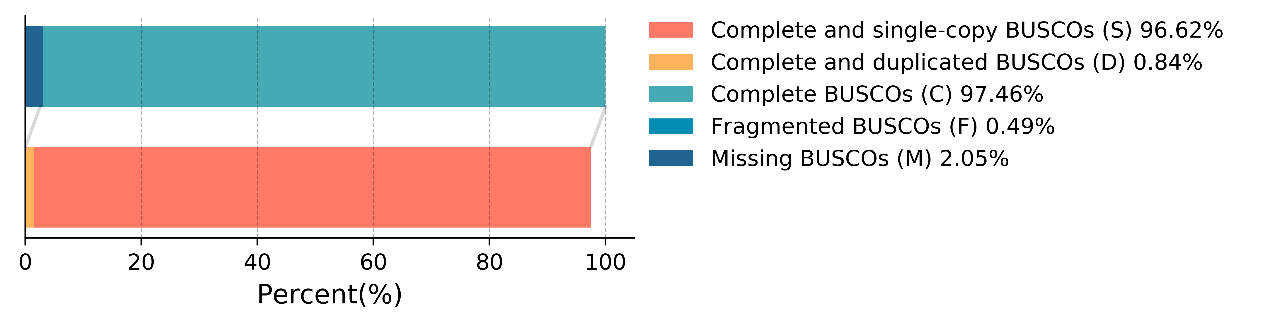

**Supplementary Figure S1 BUSCO statistics for Gnig-Y8 genome**

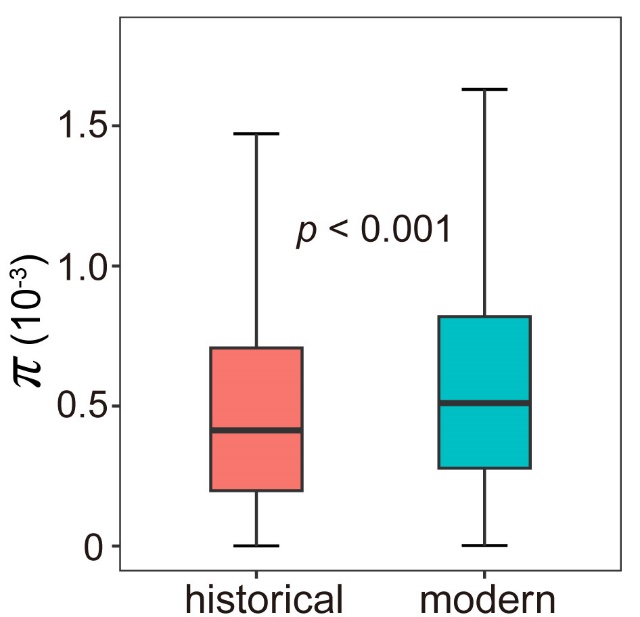

**Supplementary Figure S2 Comparison of nucleotide diversity (π) between museum and modern samples**

Boxplots comprise 1st and 3rd [quartile](https://en.wikipedia.org/wiki/Quartile)s (box), [median](https://en.wikipedia.org/wiki/Median) (line inside the box), and 1.5× interquartile range.

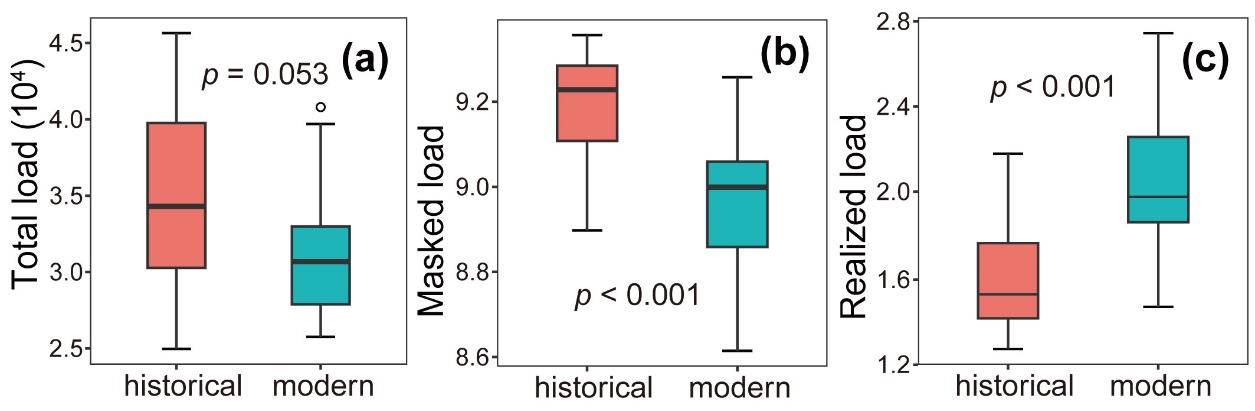

**Supplementary Figure S3 Genetic load estimation on coding regions for black-necked crane**

a: Total genetic load; b: Masked genetic load; and c: Realized genetic load based on Moderate-impact category from SnpEff analysis. Boxplot comprises 1st and 3rd [quartile](https://en.wikipedia.org/wiki/Quartile)s (box), [median](https://en.wikipedia.org/wiki/Median) (line inside box), outliers (open circle), and 1.5× interquartile range.

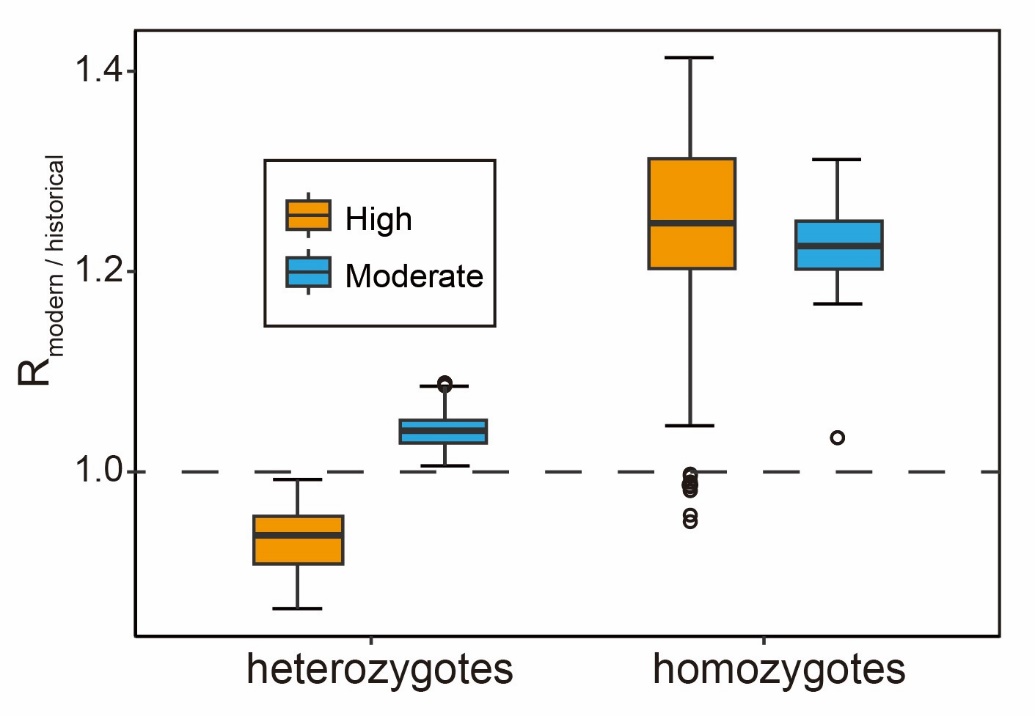

**Supplementary Figure S4 Relative frequency of derived alleles for black-necked crane**

Statistics were respectively performed for all sites (a), heterozygotes (b), and homozygotes (c) of High- and Moderate-impact variants between historical and modern genomes. Boxplot comprises 1st and 3rd [quartile](https://en.wikipedia.org/wiki/Quartile)s (box), [median](https://en.wikipedia.org/wiki/Median) (line inside box), outliers (open circle), and 1.5× interquartile range. Dashed line indicates R_modern/historical_ equal to 1, corresponding to no change in frequency between historical and modern genomes, while R_modern/historical_<1 or >1 indicates an increase or a decrease in frequency in historical relative to modern genomes, respectively. Distributions in R_modern/historical_ variance were estimated based on jack-knifing across chromosomes.

**Supplementary Table S1 Sampling and sequencing information for resequencing genomes of black-necked crane in the present study**

| Individual ID | Sampling site | Sampling time | Sampling latitude | Sampling longitude | Average coverage |
| --- | --- | --- | --- | --- | --- |
| IOZ_58199 | Haixi, Qinghai | 1959 | 37.39439022 | 97.38821644 | 33.191 |
| IOZ_58511 | Ali, Xizang | 1976 | 33.38523038 | 79.73144775 | 33.150 |
| IOZ_58512 | Ali, Xizang | 1976 | 30.29015420 | 81.17715840 | 26.858 |
| IOZ_58513 | Rikaze, Xizang | 1975 | 29.40216095 | 86.73522708 | 23.511 |
| IOZ_40401 | Xining, Qinghai | 1959 | 36.98982442 | 99.85437263 | 59.209 |
| IOZ_58514 | Ali, Xizang | 1976 | 30.29015420 | 81.17715840 | 24.520 |
| NIP_64011 | Haixi, Qinghai | 1964 | 36.94128201 | 98.47988323 | 29.498 |
| NIP_0020 | Yushu, Qinghai | 1964 | 32.99293530 | 97.00918413 | 38.266 |
| NIP_71130 | Guoluo, Qinghai | 1971 | 33.42947641 | 101.4828402 | 33.320 |
| NIP_8005 | Xining, Qinghai | 1980 | 37.56542188 | 101.1799581 | 83.815 |
| NIP_9 | Xining, Qinghai | 1980 | 37.56542188 | 101.1799581 | 32.044 |
| KIZ_20150112 | Qujing, Yunnan | 2015 | 26.69735472 | 103.2778956 | 25.323 |
| KIZ_2006120701 | Diqing, Yunnan | 2006 | 27.80594466 | 99.66145445 | 25.224 |
| KIZ_2007031602 | Diqing, Yunnan | 2007 | 27.80594466 | 99.66145445 | 25.477 |
| KIZ_2010031601 | Zhaotong, Yunnan | 2010 | 27.43587384 | 103.329344 | 23.996 |
| KIZ_2010031602 | Zhaotong, Yunnan | 2010 | 27.43587384 | 103.329344 | 23.955 |
| KIZ_2010031603 | Zhaotong, Yunnan | 2010 | 27.43587384 | 103.329344 | 25.889 |
| KIZ_2015031401 | Zhaotong, Yunnan | 2015 | 27.43587384 | 103.329344 | 24.691 |
| KIZ_2015031402 | Zhaotong, Yunnan | 2015 | 27.43587384 | 103.329344 | 25.638 |
| KIZ_2015031403 | Zhaotong, Yunnan | 2015 | 27.43587384 | 103.329344 | 25.103 |
| KIZ_20161117LU | Zhaotong, Yunnan | 2016 | 27.64028050 | 103.6539416 | 25.682 |
| KIZ_ALC47 | Ali, Xizang | 2018 | 33.56972222 | 79.94611111 | 51.356 |
| KIZ_ALC50 | Ali, Xizang | 2018 | 33.56972222 | 79.94611111 | 25.498 |
| KIZ_ALC57 | Ali, Xizang | 2018 | 33.56972222 | 79.94611111 | 37.277 |
| KIZ_ALC62 | Ali, Xizang | 2018 | 33.56972222 | 79.94611111 | 43.571 |
| KIZ_ALC7 | Ali, Xizang | 2018 | 33.56972222 | 79.94611111 | 59.532 |
| KIZ_crane_H | Zhaotong, Yunnan | 2009 | 27.43587384 | 103.329344 | 24.744 |
| KIZ_crane_J | Diqing, Yunnan | 2010 | 27.43587384 | 103.329344 | 24.997 |
| KIZ_crane_K | Diqing, Yunnan | 2010 | 27.64028050 | 103.6539416 | 25.020 |
| KIZ_crane_O | Zhaotong, Yunnan | 2010 | 27.43587384 | 103.329344 | 24.999 |
| KIZ_crane_Q | Zhaotong, Yunnan | 2009 | 27.43587384 | 103.329344 | 25.361 |
| KIZ_crane_U | Zhaotong, Yunnan | 2010 | 27.43587384 | 103.329344 | 24.887 |
| KIZ_DSB_A | Zhaotong, Yunnan | 2009 | 27.43587384 | 103.329344 | 26.134 |
| KIZ_DSB0311A | Zhaotong, Yunnan | 2010 | 27.43587384 | 103.329344 | 25.240 |
| KIZ_DSB0311B | Zhaotong, Yunnan | 2010 | 27.43587384 | 103.329344 | 25.009 |
| KIZ_NP005 | Diqing, Yunnan | 2009 | 27.80594466 | 99.66145445 | 25.879 |
| KIZ_NP008 | Diqing, Yunnan | 2010 | 27.80594466 | 99.66145445 | 25.113 |
| KIZ_NP011 | Diqing, Yunnan | 2009 | 27.80594466 | 99.66145445 | 25.326 |
| KIZ_NPH_B | Diqing, Yunnan | 2010 | 27.80594466 | 99.66145445 | 26.081 |
| KIZ_NPH090314 | Diqing, Yunnan | 2009 | 27.80594466 | 99.66145445 | 23.133 |
| KIZ_NPH200903 | Diqing, Yunnan | 2009 | 27.80594466 | 99.66145445 | 24.480 |
| KIZ_XZ3 | Ali, Xizang | 2017 | 33.20166667 | 79.8175 | 67.948 |
| TPI_YL202201 | Lasa, Xizang | 2022 | 29.91951467 | 91.09068612 | 15.857 |
| TPI_YL202208 | Lasa, Xizang | 2022 | 29.91951467 | 91.09068612 | 23.922 |
| LZU_ZLX_01 | Jiuquan,Gansu | 2020 | 40.34345641 | 93.75734047 | 25.505 |
| LZU_ZLX_02 | Jiuquan,Gansu | 2020 | 41.34345641 | 94.75734047 | 24.634 |
| LZU_ZLX_R14 | Aba, Sichuan | 2020 | 33.925628 | 102.815487 | 25.339 |
| LZU_ZLX_R18 | Aba, Sichuan | 2020 | 33.627235 | 102.684174 | 25.264 |
| LZU_ZLX_R20 | Aba, Sichuan | 2020 | 33.627235 | 102.684174 | 24.618 |
| LZU_ZLX_R22 | Aba, Sichuan | 2020 | 33.627235 | 102.684174 | 25.458 |
| LZU_ZLX_R23 | Aba, Sichuan | 2020 | 33.627235 | 102.684174 | 24.866 |
| LZU_ZLX_U00 | Gannan, Gansu | 2020 | 34.272887 | 102.305535 | 25.221 |
